# Differentiation state and culture conditions impact neural stem/progenitor cell-derived extracellular vesicle bioactivity

**DOI:** 10.1101/2023.02.15.528366

**Authors:** Dipankar Dutta, Nicholas H Pirolli, Daniel Levy, Jeffrey Tsao, Nicholas Seecharan, Xiang Xu, Zihui Wang, Xiaofeng Jia, Steven M. Jay

## Abstract

Extracellular vesicles (EVs) derived from neural progenitor/stem cells (NPSCs) have shown promising efficacy in a variety of preclinical models. However, NPSCs lack critical neuroregenerative functionality such as myelinating capacity. Further, culture conditions used in NPSC EV production lack standardization and identification of optimal conditions for NPSC EV neurogenic bioactivity. Here, we assessed whether further differentiated oligodendrocyte precursor cells (OPCs) and immature oligodendrocytes (iOLs) that give rise to mature myelinating oligodendrocytes could yield EVs with neurotherapeutic properties comparable or superior to those from NPSCs as well as mesenchymal stromal cells (MSCs), as MSC EVs are also commonly reported to have neurotherapeutic activity. We additionally examined the effects of four different extracellular matrix (ECM) coating materials (laminin, fibronectin, Matrigel, and collagen IV) and the presence or absence of growth factors (EGF, bFGF, and NGF) in cell culture on the ultimate properties of EVs. The data show that OPC EVs and iOL EVs performed similarly to NPSC EVs in PC-12 proliferation and RAW264.7 mouse macrophage antiinflammatory assays, but NPSC EVs performed better in a PC-12 neurite outgrowth assay. Additionally, the presence of nerve growth factor (NGF) in culture was found to be maximize NPSC EV bioactivity among the conditions tested. NPSC EVs produced under rationally-selected culture conditions (fibronectin + NGF) enhanced axonal regeneration and muscle reinnervation in a rat nerve crush injury model. These results highlight the impact of culture conditions on NPSC EV neuroregenerative bioactivity, thus providing additional rationale for standardization and optimization of culture conditions for NPSC EV production.

**Table of Contents:** 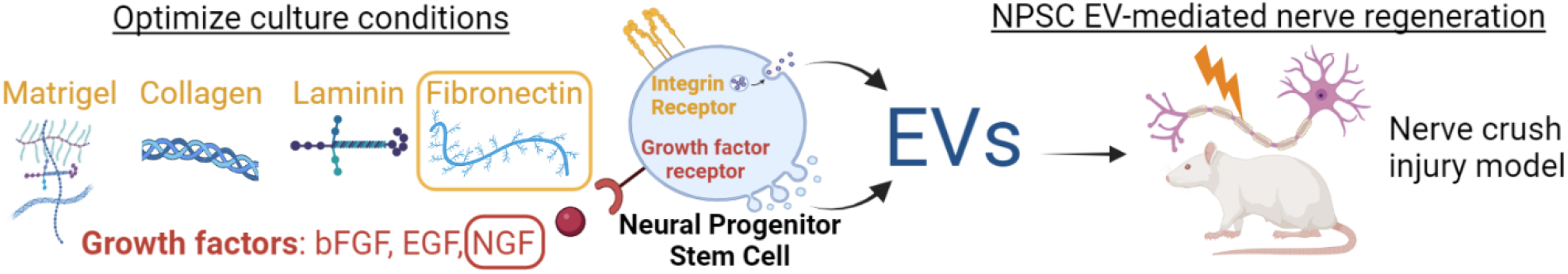

Extracellular vesicles (EVs) purified from neural progenitor/stem cells (NPSCs) have been investigated for neurotherapeutic activity, however significant variability in culture conditions limits reproducibility and efficacy of this approach. Here, we examined the impact of extracellular matrix (ECM) components and growth factors in NPSC culture on the bioactivity on the bioactivity of their EVs. The results show that EVs from NPSCs cultured with a rationally-selected ECM type (fibronectin) and growth factor (nerve growth factor (NGF)) enhanced nerve regeneration and muscle recovery in a rat sciatic nerve crush injury model.

## 1. Introduction

Extracellular vesicles (EVs) have emerged as alternatives to cell therapies for treatment of a wide variety of diseases and injuries, including spinal cord injury (SCI), peripheral nerve injury (PNI), and other neurological applications ^1,2^. While much of the early work in this area involved EVs from non-CNS source cells, especially mesenchymal stem/stromal cells (MSCs), more recent investigations into the neurotherapeutic properties of EVs from cells such as neural progenitor/stem cells (NPSCs) have yielded promising results in a variety of preclinical models ^3-5^. However, there is broad heterogeneity in how NPSCs are cultured, and culture conditions are critical to EV production and bioactivity. Thus, this lack of standardization could contribute to decreased potency of some NPSC EV formulations as well as reduced reproducibility of results within the field, both of which are limiting to the ultimate translation of EVs to treat central nervous system (CNS) diseases and injuries ^6,7^.

A critical aspect of cell culture conditions for NPSCs is the extracellular matrix (ECM) component. Stem cells such as NPSCs possess a wide variety of integrin receptors enabling cellular phenotypic responses to extracellular ECM signals, and this ECM signaling has been leveraged in vitro by culturing NPSCs on different ECM types to enhance NPSC proliferation and control their differentiation. However, it remains relatively unknown how ECM signaling can specifically influence the bioactivity of EVs released from NPSCs in culture. Additionally, the impact of supplementing different growth factors in culture media on NPSC EV bioactivity has also not been systematically explored.

Beyond culture conditions, it is notable that while NPSC EVs have received increasing attention, the neuroregenerative potential of EVs derived from oligodendrocyte precursor cells (OPCs) and their differentiated oligodendrocyte (OL) phenotypes are not well characterized, despite these latter cell types being critical to the process of myelination in the CNS. Similar to MSCs, exogenous OPC transplants correspond with functional improvements after SCI ^8-11^. Known OPC/OL-mediated paracrine effects include neuroprotection/axonal sparing ^12-15^, adaptive neural plasticity ^16^, angiogenesis ^17^, axonal growth ^18^, and endogenous oligodendrogenesis from ependymal cells ^19^. However, primary OPCs are not easily cultivated or cultured, hindering their potential as EV sources.

In this study, we addressed the issues identified above by establishing cultures of induced-pluripotent stem cell (iPSC)-derived NPSCs as well as OPCs and immature OLs (iOLs) for EV production. We observed that only NPSC EVs promoted neurite outgrowth in a PC-12 neural cell line, and that this bioactivity was modulated by both the ECM coating in adherent cell culture conditions and the presence of growth factors during EV biogenesis. Specially, we report that NPSCs cultured on laminin and fibronectin produced EVs with enhanced PC-12 neurite outgrowth compared to Matrigel and collagen IV. Additionally, mitogen withdrawal abrogated NPSC EV bioactivity across several in vitro assays, whereas supplementation with nerve growth factor (NGF) in culture generally enhanced EV bioactivity. Thus, overall we show that ECM and growth factors in NPSC culture are critical determinants of NPSC EV bioactivity and should be carefully considered and reported on in future NPSC EV studies. Standardized use of optimal ECM and culture conditions holds promise to enhance NPSC EV efficacy for a wide range of CNS therapies, enhancing their translational potential.

## 2. Experimental Section

### 2.1 Cell culture

Human induced-pluripotent stem cell (iPSC)-derived NPSCs were purchased from a commercial source (Tempo Biosciences, Tempo-iNSTem). Upon receipt, NPSCs were plated in Matrigel-coated flasks at 5,700cells/cm^2^ and labeled as passage 1 (P1). Growth factor-reduced Matrigel (Corning, 356231) was mixed in DMEM:F12 (Thermo Fisher Scientific, 11330032) under chilled conditions and added in tissue culture flasks at 60μg/cm^2^. The flasks were then incubated at 37°C for at least 1h before the Matrigel coating solution was removed to subsequently plate the cells. NPSCs were cultured in basal media that we will refer to as N2B27. N2B27 formulation is as follows: DMEM:F12 supplemented with 1X non-essential amino acids (NEAA; Thermo Fisher Scientific, 11140050), 1X Glutamax (Thermo Fisher Scientific, 35050061), 1X penicillin/streptomycin (Sigma-Aldrich, P0781), 0.1mM β-mercaptoethanol, 2μg/ml heparin, 1X N2 supplement (Thermo Fisher Scientific, 17502048), and 1X B27 supplement (Thermo Fisher Scientific, 12587010). NPSCs were expanded using NPSC proliferation media (NPM) composed of N2B27 supplemented with 20 ng/ml epidermal growth factor (EGF; Peprotech, AF-100-15), 20ng/ml basic fibroblast growth factor (bFGF; Peprotech, 100-18B) and 25μg/ml insulin (Sigma-Aldrich, I9278). NPSCs were passaged upon reaching 85-90% confluency using Accutase (Thermo fisher, A1110501). If applicable, NPSCs were stored in liquid nitrogen after suspending in NPM supplemented with 10μM Y-27632 (ROCKi; Tocris, 1254) and 1X chemically defined freezing solution (Lonza, 12-769E). 10μM ROCKi was supplemented in NPM for the first 24h after every passage or plating of frozen stocks.

Human iPSC-NPSC-derived oligodendrocyte precursor cells (OPCs) were acquired from Tempo Biosciences (Tempo-iOligo), upon receipt cells were plated at 10,000cells/cm^2^ on Matrigel-coated culture flasks and labeled as P1. OPCs were cultured using OPC maintenance/proliferation media (OMM) composed of N2B27 supplemented with 10ng/ml insulin-like growth factor (IGF-1; Peprotech, 100-11), 15ng/ml platelet-derived growth factor AA (PDGFaa; Peprotech, 100-13A), 5ng/ml hepatocyte growth factor (HGF; Peprotech, 100-39H), 10ng/ml Neurotrophin-3 (NT3; Peprotech, 450-03), 25μg/ml insulin, 1μM N6,2′-O-Dibutyryladenosine 3′,5′-cyclic monophosphate sodium salt (cAMP; Sigma-Aldrich, D0260), 100ng/ml biotin (Sigma-Aldrich, B4639) and 10ng/ml 3,3’,5-triiodo-L-thyronine sodium salt hormone (T3; Sigma-Aldrich, T6397). OPCs were passaged at 75% confluency where P2 and P3 cells were plated at 7,500cells/cm^2^. OPCs were stored in liquid nitrogen after suspending in OMM supplemented with 10μM ROCKi and 1X chemically defined freezing solution. 10μM ROCKi was supplemented in OMM for the first 24h after every passage or plating of frozen stocks. Cell populations likely to contain higher concentrations of immature oligodendrocytes (iOLs) were generated via growth factor withdrawal and exposure to high levels of triiodothyronine (T3) hormone for 3d, as described by others ^20-22^. In short, OPCs were expanded until ∼55% confluency and the media was changed to OPC differentiation media (ODM) composed of N2B27 supplemented with 1_μ_M cAMP, 25_μ_g/ml insulin and 200ng/ml T3. The OPCs were incubated for 24h and the media was subsequently changed to ODM supplemented with 4_μ_g/ml mitomycin-C (MMC; Sigma-Aldrich, M4287) overnight to arrest cell proliferation. Finally, the media was changed once more to fresh ODM without MMC for the subsequent three days.

Adipose-derived and bone marrow-derived mesenchymal stem cells (ADMSC and BDMSC, respectively) were provided by RoosterBio. The MSC cell lines were all primary human-derived with separate donors for each tissue source labeled as AD061, AD088 & AD097 for ADMSCs and BM174, BM180 & BM182 for BDMSCs. All MSCs were plated at 3000cells/cm^2^ in tissue culture-treated flasks and cultured in MSC media, consisting of Dulbecco’s Modified Eagle’s Medium (DMEM), supplemented with 10% (v/v) heat-inactivated fetal bovine serum (Hi-FBS), 1X NEAA and 1X penicillin/streptomycin. After initial plating, each cell line was labeled as P1. MSCs were passaged upon reaching 90-95% confluency and if applicable, stored in liquid nitrogen frozen in MSC media supplemented with 10% (v/v) DMSO.

### 2.2 EV isolation

P3 cells were used for EVs isolation from all cell types (NPSCs, OPCs, iOLs and MSCs), unless otherwise stated (Table 2). EV-depleted Hi-FBS was used when necessary for EV isolation and *in vitro* assays. Here, Hi-FBS was centrifuged at 100,000 x *g* for 16h and passing the top 70% of the supernatant through a 0.2μm filter before storage at -20°C. For the neural cell types (NPSCs, OPCs and iOLs), basal N2B27 underwent the same EV-depletion process as Hi-FBS before starting EV collection. Suspension (neurosphere) culture for NPSCs were initiated by suspending at 20,000cells/ml in NPM. Media was changed every 2-3d by gently pelleting the resulting neurospheres and replacing the supernatant with fresh NPM. Neurosphere diameter was visually inspected, and EV collection period was initiated when neurospheres reached a diameter of ∼0.25mm. Adherent culture conditions for EV collection included ECM signaling cues from Matrigel, fibronectin (Sigma-Aldrich, F0895-5MG), laminin (Sigma-Aldrich, L2020-1MG) and collagen IV (Sigma-Aldrich, C5533-5MG). Tissue culture flasks were incubated with fibronectin at 1.5μg/cm^2^ diluted in 1X HBSS for at least 1h at 37°C. Fibronectin coating solution was then aspirated and allowed to air-dry for 30min before plating cells. Laminin immobilization on culture flasks was completed in two steps. First, the flasks were coated with poly-L-ornithine hydrobromide (PLO; Sigma-Aldrich P4957-50ML) at 15μg/ml diluted in 1X PBS for at least 1h at 37°C. Second, laminin was diluted in DMEM:F12 to final concentration of 20μg/ml and incubated at 37°C for at least 2h. The plates were rinsed once with 1X PBS before plating cells. Collagen IV stocks were diluted in sterile dH2O for a coating concentration of 5μg/cm^2^. Plates were then coated overnight at 4°C, followed by a 1h incubation in 37°C and cells were plated after washing the coated surface once with 1X PBS.

For NPSCs and MSCs, EV isolation was initiated upon EV parent cells reaching a confluency of ∼75%. For OPCs, EV collection was initiated at ∼65% confluency (Figure 1). EV isolation for the iOLs were initiated after the 3d ODM treatment (Figure 1). For all cases, EV parent cells were incubated with EV-depleted media (where all growth factor and small molecule supplements were added immediately before use) for 48h before the conditioned media underwent a standardized EV isolation process via differential centrifugation.^23^ In short, conditioned media was centrifuged at 1000 x *g* for 10min. The supernatant was then centrifuged at 2000 x *g* for 20min and the process was repeated once more at 10,000 x *g* for 30min. Finally, small EVs were pelleted by centrifuging at 100,000 x *g* for 2h. The supernatant was carefully discarded, the pellet was resuspended in PBS. The resuspended EV-rich solutions were washed and concentrated with 1X PBS using Nanosep Centrifugal Devices with 300kDa molecular weight cut-off (Pall, OD300C35). The isolated EVs were then resuspended in 1X PBS and aliquoted for storage at -20°C for up to 8wk where the samples were subjected to no more than 1 freeze/thaw cycle.

**Figure 1.**
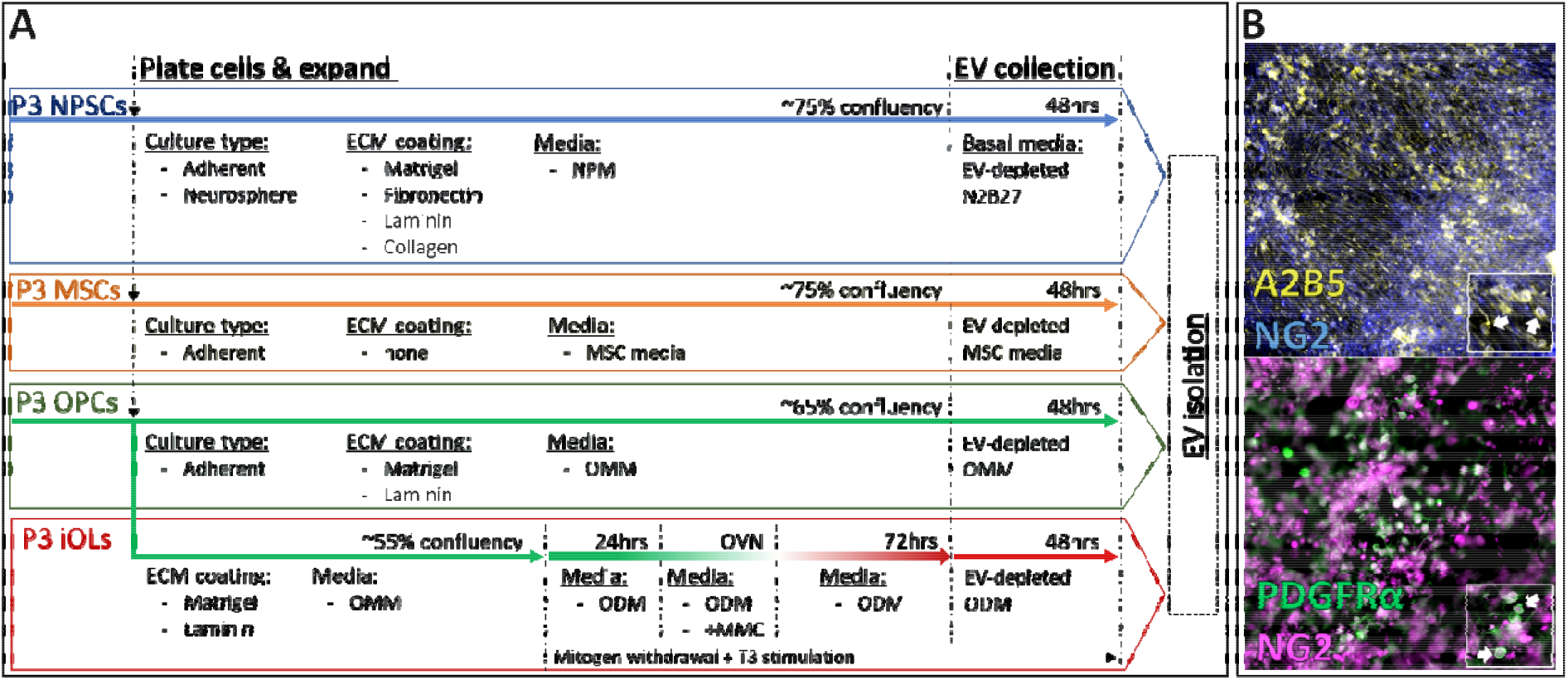
Schematic depicting all EV groups, EV isolation workflow, and OPC characterization. A) Cells were grown in the indicated culture conditions until the appropriate confluency was reached, at which point EV collection was commenced for 48h. B) Immunocytochemistry reveals presence of OPC markers A2B5 and NG2 (upper panel) and iOL markers PDGFR_α_ and NG2 (lower panel).

### 2.3 EV characterization

Protein content in isolated EV samples was determined using bicinchoninic acid assay (BCA) following the manufacturer’s protocol. Particle size and concentration were characterized via nanoparticle tracking analysis (NTA) using NanoSight (Malvern Pananalytical, LM10). In short, EV solutions were diluted to a concentration that would produce between 60-120 of particles per frame. A minimum of three 30s videos were captured using NTA analytical software version 2.3 with camera level set at 14. The detection threshold for image analysis was set at the beginning of each sample and kept constant for each replicate. Data were corrected for dilutions and presented as particle concentration (particles/ml) and percent population as a function of diameter.

#### Transmission electron microscopy

(TEM) was performed after negative staining of EVs to verify appropriate EV morphology. Briefly, EV isolates were fixed 1:1 with 4% paraformaldehyde (total volume = 40_μ_L) for 30min, and the mixture was then placed on parafilm forming a droplet.

An ultrathin carbon film grid (Electron Microscopy Sciences (Hatfield, PA)) was carefully floated on the droplet and allowed to incubate for 20min. Then, the grid was blotted to remove excess liquid, washed with PBS, and floated on 1% glutaraldehyde (50 μL, 5min) followed by dH20 washing and negative staining with uranyl acetate replacement stain (50μL, 10min). Air-dried grids were imaged with a JEM-2100 transmission electron microscope (JEOL, Japan) at 200kV accelerating voltage.

#### Molecular characterization of EVs was accomplished by immunoblot

Approximately 15μg of protein from EV isolates and lysates from respective cell types was used for immunoblots. The EV markers Alix and TSG101 and the negative markers Calnexin and GAPDH were assessed in accordance with standards from the International Society for Extracellular Vesicles.^7^ Primary antibodies for ALIX (Abcam, ab186429), TSG101 (Abcam, ab125011) and Calnexin (Cell Signaling, C5C9) were used at a 1:1000 dilution. Secondary goat anti-rabbit IRDye 800CW (LICOR, 925–32210) was used at a dilution of 1:10,000. Immunoblots were then imaged using a LI-COR Odyssey CLX Imager.

### 2.4 PC-12 Neurite Outgrowth Assay

PC-12 cells were acquired from a commercial source (ATCC, CRL1721) and expanded in PC-12 proliferation media consisting of DMEM (Corning, 10-013-CV) supplemented with 15% (v/v) Hi-FBS, 1X glutamax, 1X HEPES (Thermo Fisher Scientific, 11330032), 1X NEAA and 1X penicillin/streptomycin. PC-12 cells were expanded by suspending the cells in tissue culture-treated flasks at a concentration of 2.5×10^5^cells/ml. Media was changed every 2-3d and cells were discarded after passage 9. The neurite outgrowth assays were conducted in collagen IV-coated 96-well plates. In short, PC-12 cells were plated at 5000cells/cm^2^ and cultured with PC-12 proliferation media for the first 24h. PC-12 proliferation media was then exchanged to basal PC-12 assay media consisting of DMEM supplemented with 2% (v/v) EV-depleted Hi-FBS, 1X Glutamax, 1X HEPES, 1X NEAA and 1X penicillin/streptomycin. The controls for the neurite outgrowth assay were as follows: negative – basal PC-12 assay media; positive – basal media + 100ng/ml nerve growth factor (NGF; Peprotech, 450-01); inhibitory – basal media + 100ng/ml NGF + 0.3825μM dexamethasone. Dosage response for each EV experimental group was determined by diluting isolated EVs to 1.0×10^9^ (high), 2.8×10^8^ (med) and 2.8×10^7^ (low) particles/ml as measured using NTA. All neurite outgrowth assays were conducted in triplicates (3 wells/EV dosage for each group).

After a 72h exposure to all control and experimental conditions, neurite outgrowth from PC-12 cells was quantified using light and fluorescent microscopy. In short, the PC-12 cells were fixed using 3.7% (v/v) paraformaldehyde (Santa Cruz Biotechnology, sc-203049A) in 1X PBS for 15min. After washing gently with 1X PBS, the cells were permeabilized using 0.35% (v/v) Triton X-100 diluted in 1X PBS for 15min. After another set of gentle washes and 4′,6-diamidino-2-phenylindole (DAPI) nuclear staining (Thermo Scientific, R37606; following manufacturer’s protocol), neurite outgrowth was recorded by imaging phase contrast overlaid with DAPI channel fluorescent images of at least three distinct regions of each well. Acquired images were quantified using FIJI and the NeuronJ plugin.^24^ The DAPI fluorescent channels were used to estimate the number of cells in each field using FIJI’s particle count algorithm. The NeuronJ plugin was used to manually trace all cellular processes from the respective phase contrast images. Traces <10 μm in length were discarded and the cumulative length of >10 μm neurites was normalized by the number of nuclei detected.

### 2.5 RAW Inflammatory Assay

RAW264.7 mouse macrophage (RAW) cells were acquired from a commercial source (ATCC, CRL1721) and expanded in RAW media composed of DMEM supplemented with 5% Hi-FBS and 1X penicillin/streptomycin by plating at 17,500cells/cm^2^ in tissue culture-treated flasks and passaged upon reaching ∼80% confluency. RAW cells between passages 12-15 were used for all inflammatory assays, which comprised a 24h pre-treatment, followed by a 4h co-treatment regime. In short, RAW cells were plated at 70,000cells/cm^2^ in 48-well plates and cultured overnight using RAW media. The next day, the 24h pre-treatment regime was initiated by exchanging to EV-depleted RAW assay media with the following controls: negative – assay media-only; positive – assay media-only; inhibitory – assay media + 2.5μM dexamethasone. Pre-treatment for the experimental groups comprised of assay media + EVs. The next day, media was replaced to initiate the 4h co-treatment regime where the controls were as follows: negative – assay media-only; positive – assay media + 10ng/ml lipopolysaccharide (LPS; Sigma-Aldrich, L4391); inhibitory – assay media + 10ng/ml LPS + 2.5μM dexamethasone. Co-treatment for the experimental groups comprised assay media + 10ng/ml LPS + EVs. EVs were dosed identically to the neurite outgrowth assay with three concentrations for each group (1.0×10^9^, 2.8×10^8^ and 2.8×10^7^ particles/ml as measured using NTA) during both pre- and co-treatment regimens to determine dose-dependent responses. All inflammatory assays were conducted in triplicates (3 wells/EV dosage for each group). After co-treatment, the conditioned media was stored at -80° C and the interleukin-6 (IL-6) content was quantified via enzyme-linked immunosorbent assay (R&D Systems, DY406) following the manufacturer’s protocol.

### 2.6 Immunocytochemistry

OPC phenotype was validated via immunocytochemistry (ICC; Figure 1) probing for colocalized OPC surface biomarkers of neural/glial antigen-2 (NG2), A2B5 and platelet derived growth factor receptor alpha (PDGFRα).^25^ In short, OPCs were plated on laminin-coated 96-well plates until ∼65% confluency. The cells were then fixed using 3.7% (v/v) paraformaldehyde diluted in 1X PBS for 15 mins. The wells were gently washed 3x with 1X PBS to minimize cell detachment. The wells were then blocked for 60min at room temperature (RT) using 10% (v/v) normal donkey serum and 0.05% BSA diluted in 1X PBS. After washing carefully with 1X PBS + 0.05% (v/v) Tween-20 (PBS-T), the samples were incubated overnight at 4°C with the primary antibodies for A2B5 (1:50; R&D Systems, MAB1416) and NG2 (1:300; EMD Millipore, AB5320) as well as PDGFR-α (1:25; R&D Systems AF-307) and NG2, separately. After washing with PBS-T, samples were incubated with the appropriate secondary antibodies (Life Technologies, A-21432, A-31573 & A-21202) for 1h at RT. After washing 3x with PBS-T, the samples were imaged using a fluorescence microscope. Control wells without primary antibodies added were used to check for background and non-specific binding. The PC-12 neurite outgrowth assay was initially validated by probing for β-tubulin III (1:1000; Abcam, ab7751) using ICC to optimize the positive, negative, and inhibitory controls (Figure 5a). The same protocol as above was used along with an additional permeabilization step immediately before blocking. Samples were incubated with 1X PBS with 0.35% (v/v) Triton X-100 for 15min such that the primary antibody was available for binding to cytosolic β-tubulin III.

### 2.7 In-vivo studies

#### 2.7.1 Animals and Surgery Procedures

The study was approved by the Institute Animal Care and Use Committee (IACUC) of the University of Maryland, Baltimore. All animals had free access to food and water and were maintained in accordance with NIH guidelines for the humane care of animals. In brief, a total of 8 female rats (200-250g, Charles River, Wilmington, MA, USA) were randomly assigned into 2 groups: the control group (n=4) and the NPSC-EV treatment group (n=4). All the rats received anesthesia by inhalation of isoflurane and the surgical field on the randomly selected left leg of each rat was shaved and sterilely prepared. After exposing the sciatic nerve, a crush injury was created at the middle part of the exposed nerve using a surgical hemostatic forceps for 30 seconds (a 10s click for 3 times with a 10s interval between each click). After crush injury, either EV solution (10μg, EV group) or equal volume of PBS (control group) was gradually injected into the crush injury site using a syringe under a stereomicroscope. Then, a 9-0 nylon epineural stitch was used to mark the injury site at both the proximal and distal end of the injury. After closing the muscle and skin with 4-0 nylon sutures, all rats were returned to their cages and kept for 4wk. After recovery, all rats were euthanized, and nerve and muscle were harvested for further evaluations.

#### 2.7.2 Immunofluorescence staining

At 4wk after surgery, each rat was deeply euthanized and transcardially perfused with a 4% paraformaldehyde (PFA) solution. After perfusion, the sciatic nerve sections at the injury site were harvested and fixed in 4% PFA for overnight, then treated with 30% sucrose until further immunofluorescence staining. Prior to immunohistochemistry, the sciatic nerve specimens were cut into 10μm thick sections and mounted to slides. Standard immunofluorescence staining was performed as we previously described ^26^. In brief, after blocking, the nerve sections were incubated with the following primary antibodies overnight at 4 °C: rabbit anti-Neurofilament 200 (NF-200) (1:500, MilliporeSigma) and mouse anti-MBP (1:50, Santa Cruz), and then incubated with the secondary antibodies as follows: goat anti-rabbit IgG antibodies conjugated with Alexa 488 (1:500, Invitrogen) and donkey anti-mouse IgG antibodies conjugated with Alexa 594 (1:500, Invitrogen) at RT for 1h. The images were collected from at least three sections of nerve tissues for each rat under a Leica DMi8 fluorescent microscope (Leica Microsystems). Quantitative analysis of images was performed using Image J (v1.8.0, NIH, USA). Comparable images have been equally adjusted for brightness/contrast. Axon numbers and diameters were counted at three randomized microscopic fields in each section, and averages were used for further analysis.

#### 2.7.3 Wet weight of gastrocnemius muscle

Gastrocnemius muscles were excised from both the injured and contralateral side, and immediately weighed using an electronic balance (Mettler AE 160, USA). Gastrocnemius Muscle Index (GMI) was used to evaluate the muscle reinnervation following denervation-induced muscle atrophy ^26^. GMI was calculated by dividing the muscle mass at the injured side by that at the contralateral side.

### 2.8 Statistical analyses

All analyses were performed in triplicate. Data are presented as mean□±□SEM. One-way ANOVAs with Holm-Šídák multiple comparison tests were used to determine statistical significance. All statistical analysis was performed with Prism 7 (GraphPad Software, La Jolla, CA). Notation for significance in Figures are as follows: ns = P□>□0.05, * = P□<□0.05; ** = P□<□0.01; *** = P□<□0.001.

## 3. Results

### 3.1 EVs from NPSCs, OPCs, and iOLs induce dose-dependent PC-12 proliferation

An initial goal of these studies was to assess how differentiation of NPSCs into OPCs or iOLs might affect EV bioactivity. We first confirmed EV identity for each population via assessment of particle size, protein markers and vesicle morphology from these cells under various culture conditions. EVs from BDMSCs and ADMSCs were utilized as controls in various bioactivity assays and thus were also characterized. EVs from all cell types displayed similar size distributions despite various ECM culture conditions (Supplemental Figure 1). As shown for NPSCs grown on Matrigel, the presence or absence of growth factors had no discernable impact on EV size distribution, production yields, or marker expression (TSG101, Alix) (Figure 2A,B, Supplemental Table 1). Transmission electron micrographs revealed 40-200nm diameter EVs with the expected cup-shaped morphology (Figure 2C).

**Table 1.** Summary of the media formulations used for each of the cell types in the study.

| Media Type | Formulation |
| --- | --- |
| Neural basal media (N2B27) | DMEM:F12, 1x N2 supplement, 1x B27 supplement, 1x non-essential amino acids, 1x glutamax, 1x penicillin/streptomycin, 0.1mM β-mercaptoethanol and 2μg/ml heparin |
| NPSC proliferation media (NPM) | N2B27, 20ng/ml EGF, 20ng/ml bFGF and 25μg/ml insulin |
| OPC maintenance/proliferation media (OMM) | N2B27, 10ng/ml IGF-1, 15ng/ml PDGF$\alpha\alpha$, 5ng/ml HGF, 10ng/ml NT3, 25μg/ml insulin, 1 μM cAMP, 100ng/ml biotin and 10ng/ml T3 |
| OPC differentiation media (ODM) | N2B27, 1 μM cAMP, 25 μg/ml insulin and 200ng/ml T3 |
| MSC media | DMEM, 10% (v/v) Hi-FBS, 1x non-essential amino acids and 1x penicillin/streptomycin |

**Figure 2.**
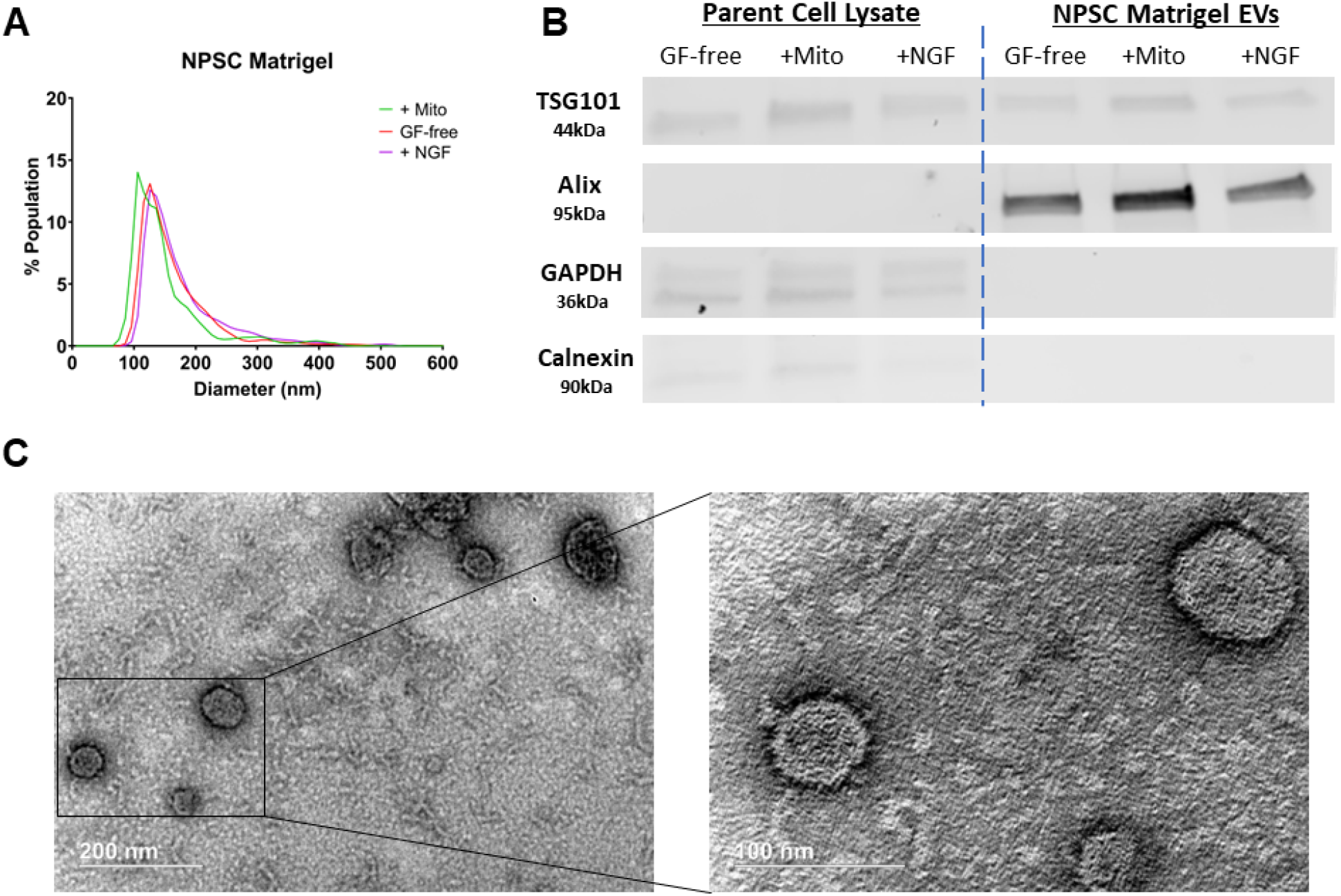
EV characterization A) Nanoparticle tracking analysis of EVs from NPSCs grown on Matrigel in the presence or absence of growth factors (EGF/bFGF) or NGF. **B)** Immunoblots of NPSC EV lysates generated from the same conditions as above for EV-associated markers TSG101 and Alix as well as negative markers calnexin and GAPDH. **C)** Transmission electron microscopy of EVs generated from NPSCs grown on Matrigel and in the presence of NGF (scale bar = 200 nm, inset 100 nm).

To assess bioactivity, a PC-12 proliferation assay was employed, as PC-12 cells are widely used as a model for neuron differentiation and neuroregeneration. MSC EVs have previously been shown to promote PC-12 cell proliferation ^27^, however, we did not observe increased PC-12 proliferation from ADMSC or BDMSC EVs across multiple donors (Figure 3A,B). NPSC EVs did show a general trend for dose-dependent increases in PC-12 proliferation across all ECM and growth factors groups, except for suspension-grown NPSC EVs, which had no effect on PC-12 proliferation (Figure 3C). Furthermore, certain combinations of ECM and growth factors produced NPSC EVs with significant dose-dependent increases in PC-12 proliferation; most notably, growth factor free (GF-free) laminin EVs produced a significant 3.5-fold increase in PC-12 proliferation. Additionally, GF-free collagen IV, and several NGF+ EV groups (fibronectin, Matrigel, collagen IV) showed significant dose-dependent increased PC-12 proliferation, but responses were variable and none reached statistical significance over basal media. These data show that the NPSC microenvironment (ECM and growth factors) can impact EV bioactivity related to PC-12 proliferation. Additionally, EVs from OPCs and iOLs cultured on laminin produced dose-dependent increases in PC-12 cell proliferation (Figure 3C). Compared to other ECMs, laminin more consistently enhanced PC-12 proliferation across multiple cell types and without a requirement for growth factors in culture media.

**Figure 3.**
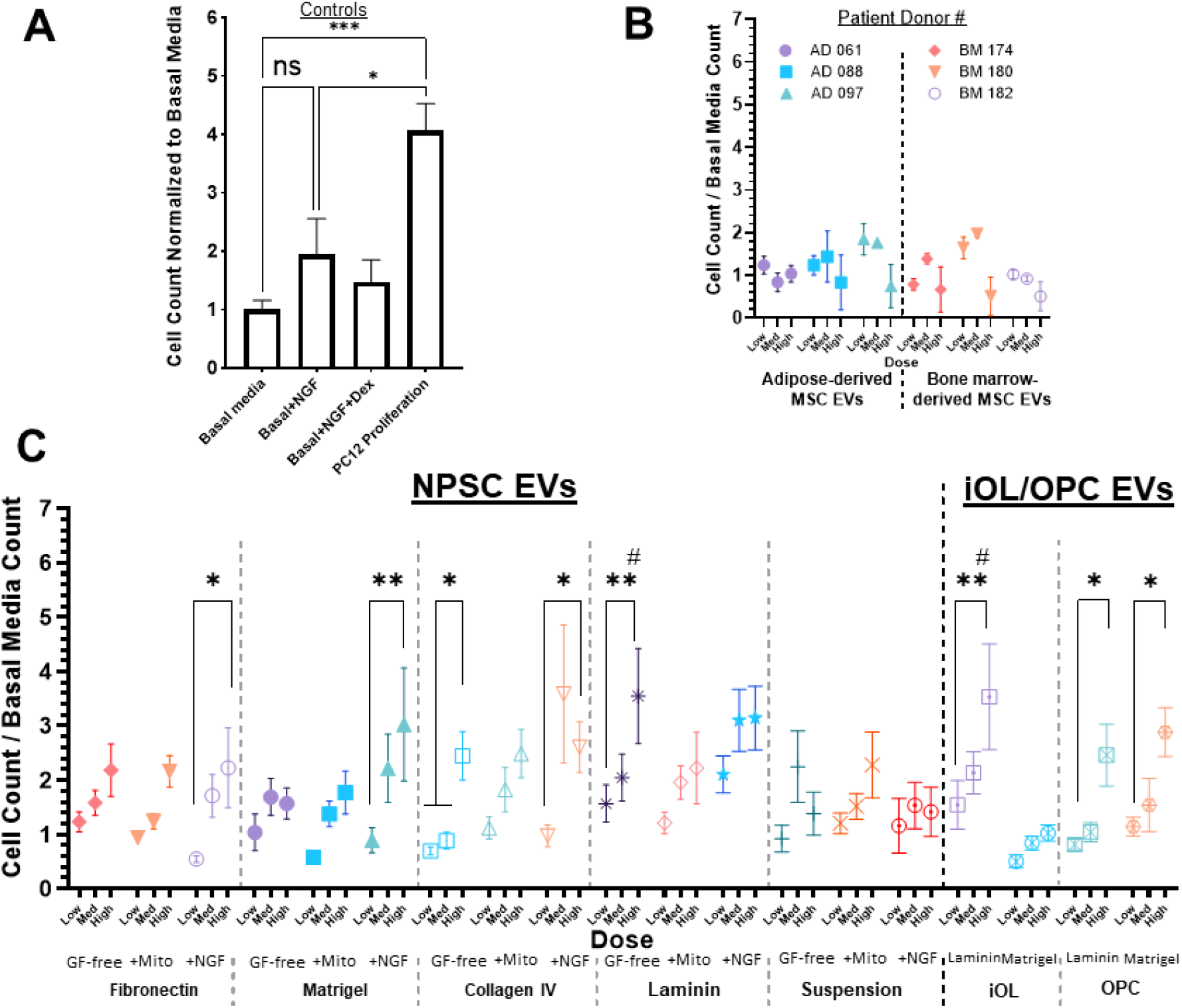
Assessment of EV-induced PC-12 proliferation. A-C) Normalized PC-12 cell count following treatment with **A)** control groups: i) basal media (negative control), ii) basal+NGF (positive control), iii) basal+NGF+Dex (inhibitory control), and iv) PC-12 proliferation media (positive control). **B)** MSC EVs from either adipose or bone marrow tissue across different patient donors, and **C)** NPSCs cultured on different ECM and in the presence of the growth factors EGF/bFGF (+Mito), or NGF (+NGF), or absence of any growth factors (GF-free), as well as OPC and iOLs cultured on either laminin or Matrigel. Doses for EV groups were 2.8e7 particles/mL (Low), 2.8e8 particles/mL (Med), and 1.0e9 particles/mL (High). All data are normalized to basal media PC-12 cell count and represented as mean ± SEM, and significance was determined with one-way (control groups) or two-way (experimental groups) ANOVA with Tukey’s post hoc test (ns: P >0.05; *P < 0.05; **P < 0.01; ****P < 0.0001 compared between doses, and # p< 0.05 compared to basal media).

### 3.2 EVs from NPSCs, OPCs, and iOLs induce dose-dependent anti-inflammatory effects in RAW264.7 macrophages

A therapeutic target for neurodegenerative disease and brain injury is reducing neuroinflammation which is primarily mediated by CNS macrophages (microglia). Thus, we sought to investigate the anti-inflammatory potential of NPSC, OPC, and iOL EVs. LPS stimulation of RAW264.7 mouse macrophages generally produced levels of the inflammatory cytokine IL-6 of >1000pg/ml in conditioned media across multiple assays (Figure 4A). Preliminary studies showed that co-treatment of EVs alone was insufficient to observe an anti-inflammatory effect (Supplemental Figure 2B). When macrophages were pre- and co-treated with EVs from all cell sources (NPSC, OPC, iOL, and MSCs as known anti-inflammatory positive controls) we observed dose dependent decreases in IL-6 levels well below 1000pg/ml indicating an anti-inflammatory effect for all cell types. EV anti-inflammatory effects were not significantly affected by presence or absence of growth factors (Figures 4B,C). Additionally, type of ECM did not have a major impact on anti-inflammatory efficacy of EVs (Figures 4D,E).

**Figure 4.**
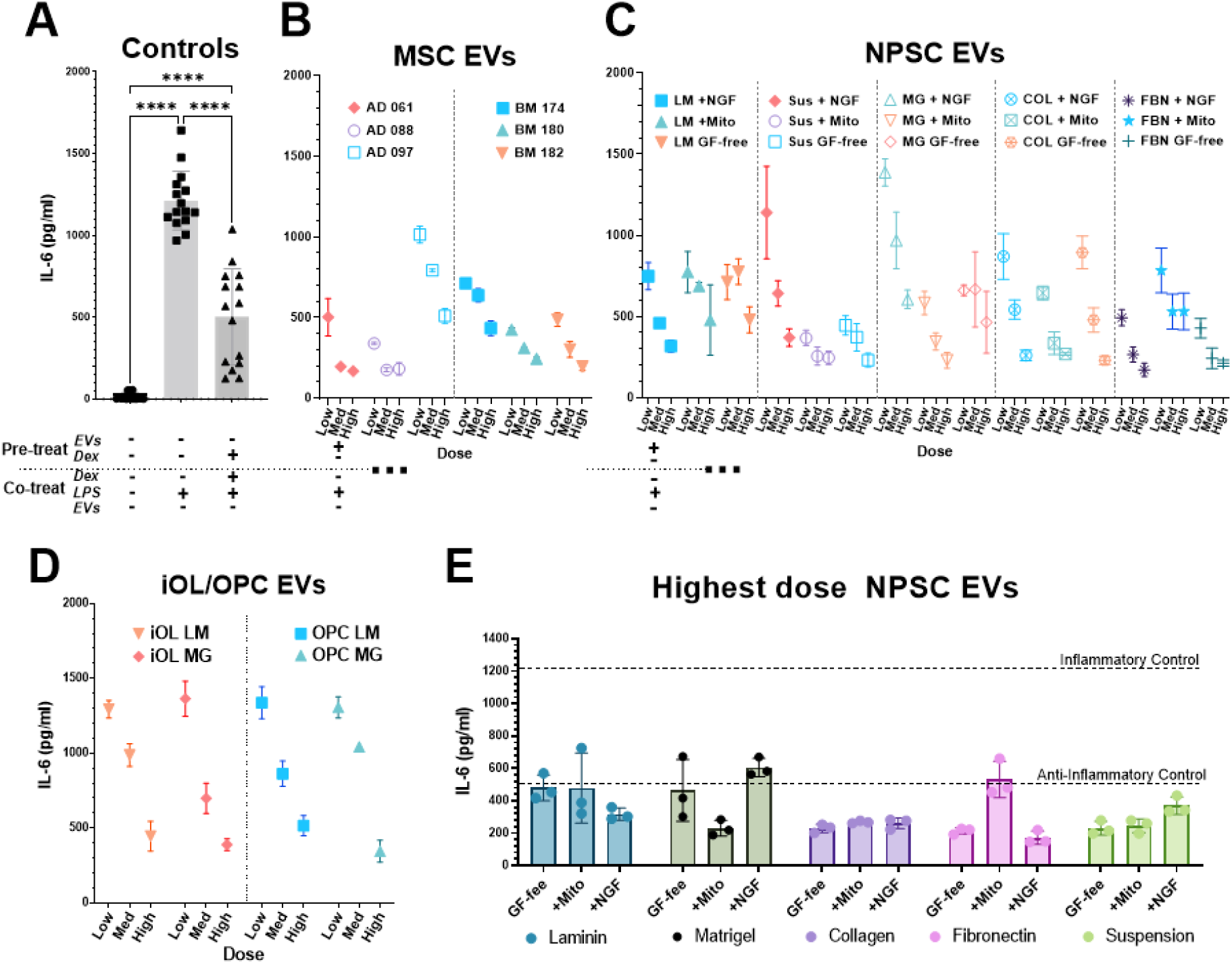
Assessment of EV anti-inflammatory bioactivity. A-D) Inflammatory responses from RAW264.7 mouse macrophages were evaluated by IL-6 levels in conditioned media following LPS-stimulation and treatment with **A)** dexamethasone (positive control), **B)** MSC EVs, and **C)** NPSC EVs, and **D)** OPC, and iOL EVs. IL-6 levels from control groups across multiple independent assays are pooled and displayed together. **E)** RAW264.7 macrophages were pre- and co-treated with EVs from NPSCs cultured in suspension or on different ECM (laminin, Matrigel, collagen IV, fibronectin), and either without growth factors (GF-free), or with EGF/bFGF (+Mito) or NGF (+NGF). IL-6 levels from inflammatory control (LPS only) and anti-inflammatory control (LPS+Dexamethasone) are indicated by dashed horizontal lines. Doses for EV groups were 2.8e7 particles/mL (Low), 2.8e8 particles/mL (Med), and 1.0e9 particles/ML (High). Statistical significance for control groups was determined by one-way ANOVA with Tukey’s post hoc test (****P < 0.0001).

### 3.3 NPSC EVs stimulate PC-12 neurite outgrowth

Next, we utilized a PC-12 neurite outgrowth assay as a more relevant *in vitro* model for neuron differentiation and growth. NGF was used a positive control benchmark and produced a cell-count normalized average sum neurite length of 12.1μm (Figure 5A) relative to the basal media control group, where the normalized neurite length was approximately 1μm. We assessed several methods for neurite outgrowth data analysis and found similar trends in control groups across all methods. Cumulative sum of neurites >10μm normalized by cell count was selected due to the relatively low standard deviations between replicates and the statistically significant differences across controls compared to other methods (Supplemental Figure 2A). This method also normalizes for cell count which can vary significantly across individual images that are being quantified. EVs from OPCs, iOLs, ADMSCs, or BDMSCs did not increase PC-12 neurite outgrowth above basal media levels (Figure 5B). However, NPSC EVs from most culture conditions showed dose-dependent increases in neurite outgrowth, reaching similar levels as the NGF positive control (Figure 5B). Overall, these results indicate that differentiation of NPSCs in to iOLs or OPCs could diminish some of the beneficial bioactivity of EVs and confirm NPSCs as a promising source for generation of EVs with neuroregenerative properties.

**Figure 5.**
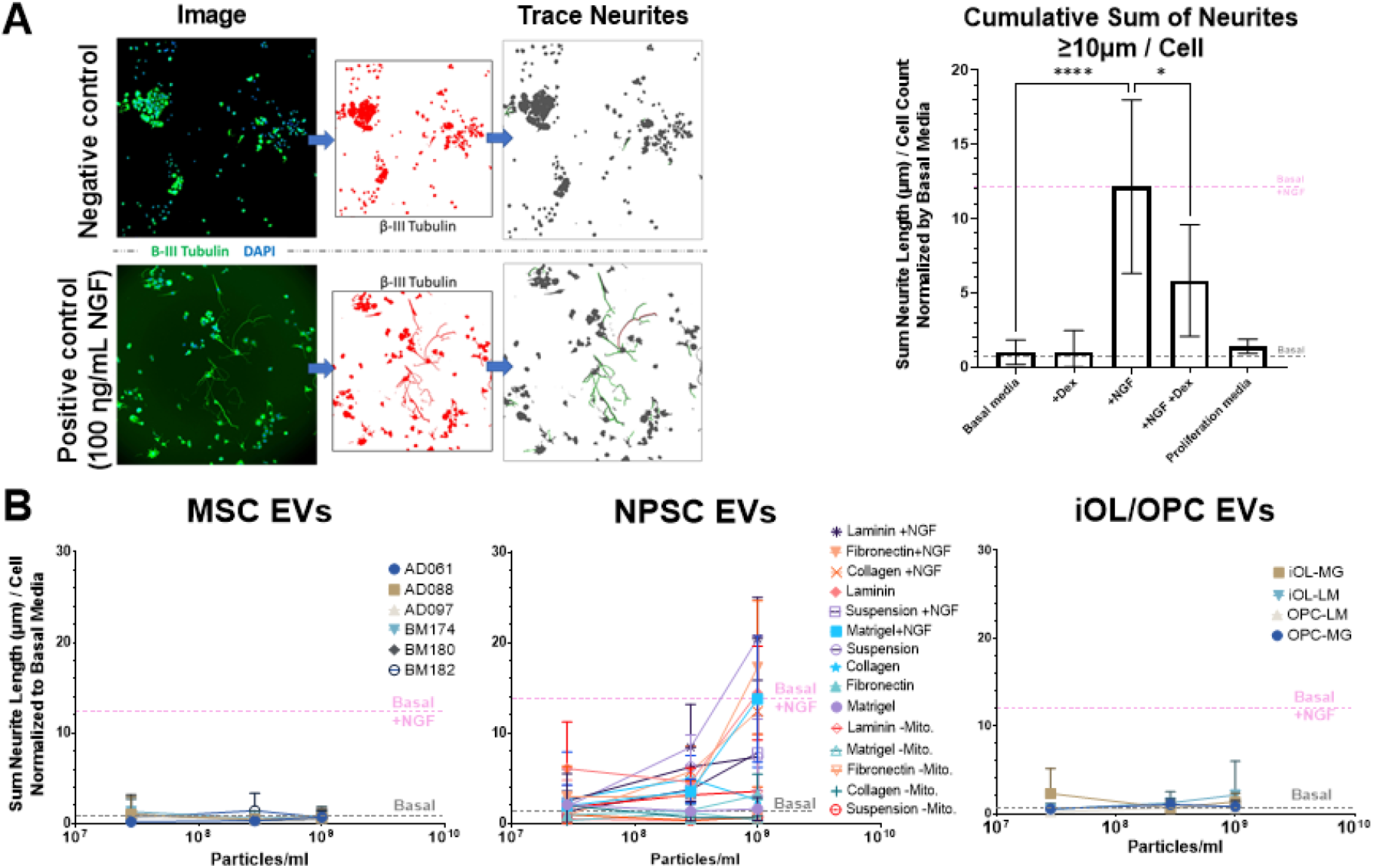
NPSC EVs stimulate neurite extension in vitro to a greater degree than OPC and iOL EVs. A) (left) β-III tubulin immunocytochemistry to label and quantify PC-12 neurite length during assay development; (right) control groups: basal media and proliferation media (negative controls), dexamethasone (+Dex) and +NGF+Dex (inhibitory controls), and +NGF (positive control). NGF produced a mean normalized PC-12 neurite length of ∼12 μm which is indicated by a pink horizontal line throughout subsequent figures. **B)** PC-12 neurite outgrowth of MSC EVs (left), NPSC EVs (middle), and OPC/iOL EVs (right) across culture conditions and dosage. Data presented as cumulative sum of all neurites greater than 10 μm normalized by cell count and basal media PC-12 neurite length. Statistical significance for control groups was determined by one-way ANOVA with Tukey’s post hoc test (*P < 0.05; ****P < 0.0001).

### 3.4 Integrin and growth factor signaling potentiate neurite outgrowth bioactivity of NPSC EVs

Having identified NPSCs as the most promising source of neurotherapeutic EVs among those we investigated, we next sought to further investigate specific culture conditions that best enhance NPSC EV PC-12 neurite outgrowth bioactivity. Among the four ECM conditions tested, NPSCs cultured on laminin produced EVs with significantly enhanced PC-12 neurite outgrowth bioactivity compared to all other ECM with average values above the NGF positive control (Figure 6A). EVs from suspension-grown NPSCs also enhanced PC-12 neurite outgrowth, but less than laminin and at levels insufficient to reach statistical significance over other ECM groups. Next, we tested the same four ECMs with and without growth factors NGF or EGF/bFGF. EGF/bFGF are typically included in NPSC culture, whereas NGF is not. NGF alone (without EGF/bFGF) significantly enhanced PC-12 neurite outgrowth across all ECMs regardless of the presence of EGF/bFGF (Figure 6B). Overall, the laminin+NGF and fibronectin+NGF groups produced the greatest PC-12 neurite outgrowth compared to all other conditions, and laminin+NGF produced statistically significant increases above the NGF positive control (Figure 6B). Thus, culturing NPSCs with specific combinations of integrins and growth factors is a potential method for enhancing NPSC EV-mediated neurite outgrowth bioactivity, with the presence of NGF being a critical factor.

**Figure 6.**
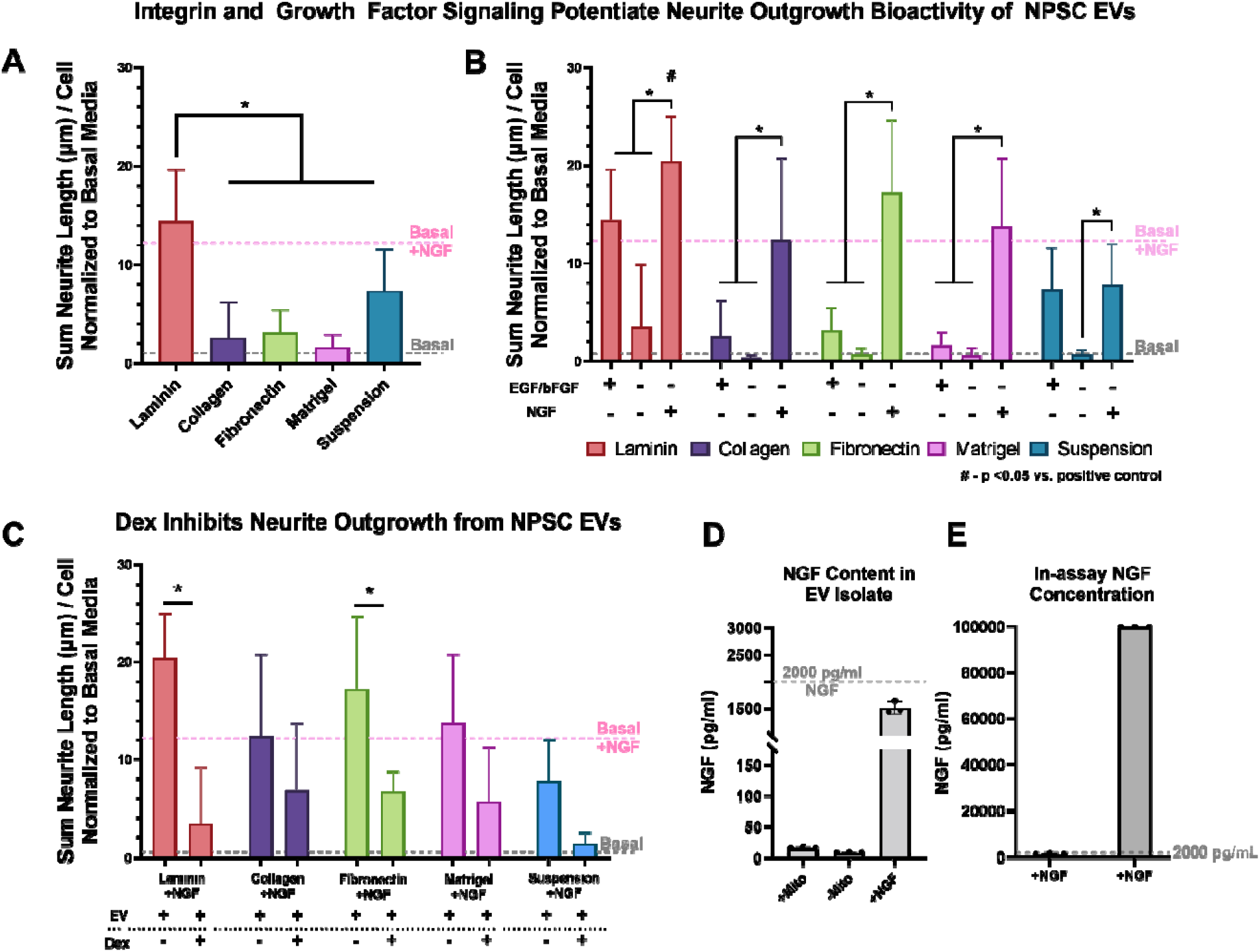
Integrin and growth factor signaling potentiate neurite outgrowth bioactivity of NPSC EVs. A-C) PC-12 neurite lengths following treatment with EVs from **A)** NPSCs growth on different ECM (laminin, collagen, fibronectin, Matrigel) or grown in suspension, **B)** NPSCs cultured with the same ECM but with EGF/bFGF, or with NGF, or without any growth factors, and **C)** co-treatment with dexamethasone in addition to EVs from NPSCs cultured with NGF. Mean neurite lengths following treatment with basal media only or basal media+NGF are indicated by dashed horizontal lines. **D-E)** Concentration of co-isolated NGF in purified EV samples (D) and calculated effective concentration in neurite outgrowth assay compared to 100 ng/ml NGF positive control (E). Data represent “High” EV dose for all groups was 1.0e9 particles/mL. Statistical significance was determined with one-way (A) or two-way (B-C) ANOVA with Tukey’s post hoc test (*P < 0.05; **P < 0.01; ****P < 0.0001 compared between dosage or groups, and # p< 0.05 compared to basal media + NGF positive control).

Dexamethasone is a glucocorticoid receptor agonist known to inhibit neurite outgrowth in PC-12 cells. In our studies, dexamethasone exhibited a trend for suppression of NPSC EV neurite outgrowth bioactivity across all culture conditions tested, with statistically significant suppression of laminin+NGF and fibronectin+NGF NPSC EV bioactivity and similar trends for the other groups tested (Figure 6C). These results suggest that NPSC EVs may increase neurite outgrowth via similar mechanisms that are suppressed by dexamethasone. Finally, since 100ng/mL NGF is a positive control for neurite outgrowth, NGF in NPSC culture media could potentially enhance EV bioactivity via co-isolation of NGF with EVs during EV purification. We found that although NGF was present in EV samples derived from NPSCs cultured with NGF, the highest effective NGF protein concentration (as measured by ELISA) during PC-12 assays was <2ng/ml, relative to the positive control (100ng/ml) (Figures 6D,E). Additionally, we tested multiple EV isolation methods (ultracentrifugation or tangential flow filtration) and found fibronectin+NGF EVs had similar PC-12 neurite outgrowth as well as anti-inflammatory bioactivity across isolation mesthods (Supplemental Figure 3). Thus, it is unlikely that co-isolated NGF solely accounts for the increased EV bioactivity above levels of the 100 ng/ml NGF control seen with laminin+NGF and fibronectin+NGF groups. However, we cannot exclude a minor contribution of co-isolated NGF to these results.

### 3.5 NPSC EVs stimulated axonal regeneration and attenuated muscle atrophy in vivo

In order to investigate the effects of EVs delivered from optimized NPSC culture conditions on axonal regeneration in a peripheral nerve crush injury model, we selected fibronectin+NGF NPSC EVs for further study based on their relative effectiveness and consistency in the *in vitro* assays. Neurofilament 200 (NF200) was used to represented neurofilaments for axonal tracing and myelin basic protein (MBP) was used as a marker for re-myelination on harvested sciatic nerve (Fig 7A). NF200 contributes substantially to axon structure and axon diameter by providing structural support, and its arrangement is parallel to axon growth and can reflect the number of axons. The results showed that at 4wk after surgery, NPSC-derived EVs significantly increased axon count and axon diameter compared to control treatment (PBS). Quantitative analysis indicated that the total number of NF200-positive axons in the NPSC EV group was significantly higher than that in the control group (302.8 ± 46.2 vs. 118.6 ± 8.0; p<0.05) (Fig 7A). In addition, the diameter of NF200-positive axons in the NPSC EV groups was 5-fold higher than that in the control group (21.4 ± 3.1 vs. 4.3 ± 1.0; p<0.001) (Fig 7A). These observations indicate that significantly more axons, with larger diameters, regenerated based on NPSC EV treatment.

**Figure 7.**
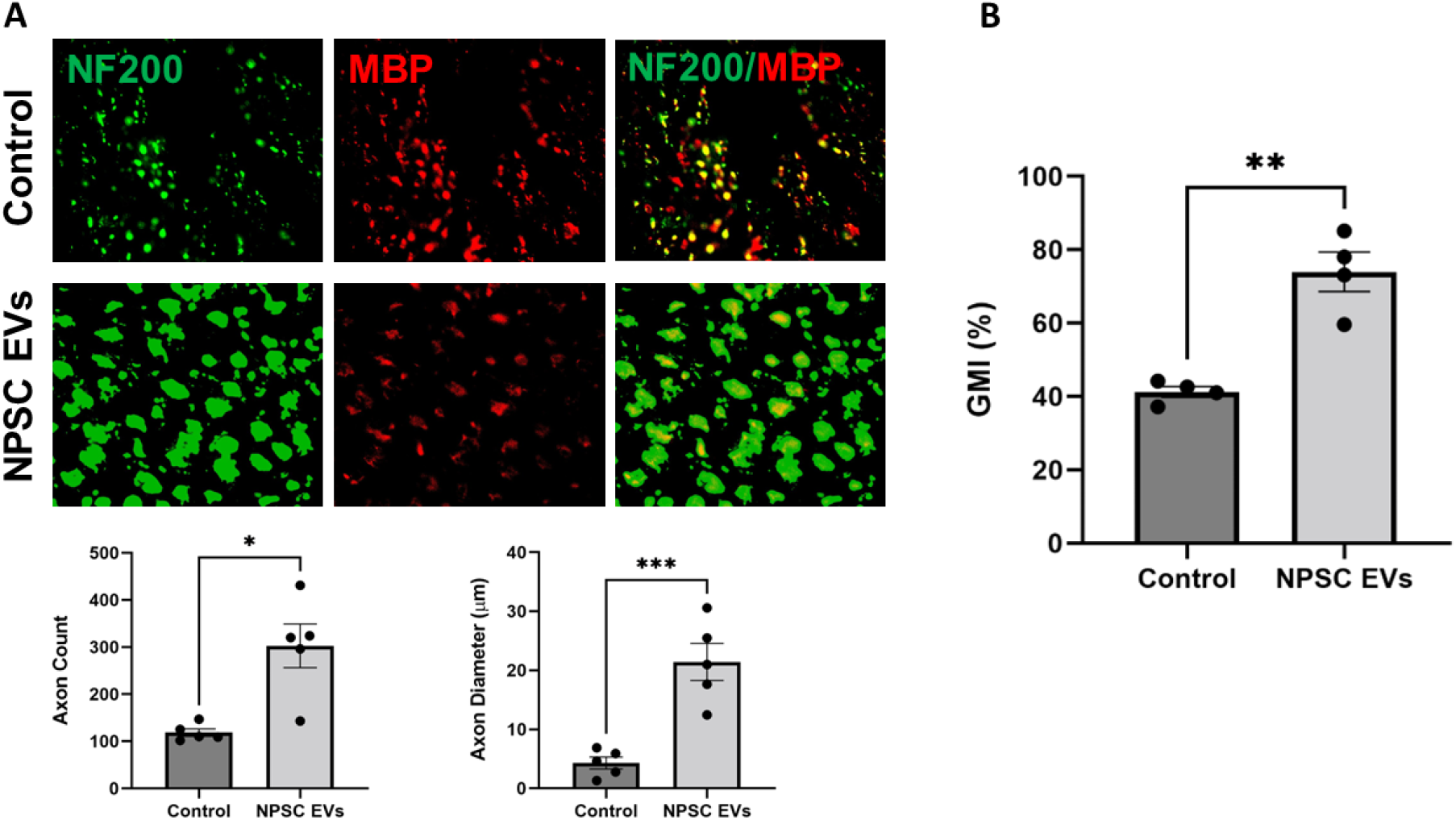
NPSC EVs promote myelinated axonal regeneration and muscle reinnervation in a rat sciatic nerve crush injury model. A) Analysis of nerve regeneration in animals treated with control (PBS) and NPSC EVs (n=4). (Top) Immunohistochemical staining for NF200 and MBP on labeled myelinated axons in cross sections at 4wk following sciatic nerve crush injury. (Bottom) Quantitative analysis of NF200-positive axons via ImageJ. **B)** Gastrocnemius wet muscle weight analysis of animals administered control (PBS) or NPSC EVs (n=4).

A further assessment of muscle recovery during nerve regeneration was conducted by measuring the wet muscle weight on the gastrocnemius muscle on the injured side of the animal and normalizing against the weight of the same muscle on the non-injured side to establish a gastrocnemius muscle index (GMI). The GMI is a standardized measurement of gastrocnemius muscle weight, which is expected to initially decrease after nerve injury due to atrophy and can subsequently partially recover based on successful reinnervation ^26^. Therefore, the GMI serves as an indirect measure of nerve regeneration. In this study, the GMI was significantly higher in the NPSC EV group compared to that of the control group (74.0 ± 5.4 vs. 41.3 ± 1.5; p<0.01) (Figure 7B), indicating that NPSC EVs attenuated muscle atrophy after crush nerve injury.

## 4. Discussion

NPSC EVs are a promising alternative to cellular therapies for neuroregenerative applications. Here, we tested the hypothesis that differentiating NPSC to OPCs and/or iOLs could improve the neuroregenerative properties of their EVs based on the myelinating capacity of the latter two cell types. Our results do not support this hypothesis, although it is important to mention that the assays used here represent only a small fraction of the analyses that could be applied to assess EV neurotherapeutic potential. Nevertheless, based on our data, we shifted focus to further investigate the potential of NPSC EVs for neurological applications. Specifically, we investigated the effects of culture conditions on NPSC EV bioactivity. Parameters such as ECM coating of cell culture substrates are well known to modulate NPSC bioactivity, but their impact on NPSC EV bioactivity had not been investigated. Laminin, Matrigel, fibronectin, and collagen IV are commonly used coating materials for NPSC culture and provide a biomimetic microenvironment and integrin signaling to promote NPSC proliferation ^29^, differentiation to neurons ^30^, viability ^31^, among other functions. In particular, prior reports have indicated that laminin produced superior NPSC proliferation and cell number compared to other ECMs ^32^. Here, we show that ECM type is also a critical determinant of NPSC EV bioactivity. NPSCs grown on laminin or fibronectin produced EVs that enhanced PC-12 neurite outgrowth and proliferation compared to Matrigel and collagen IV. Indeed, prior studies have shown *ex vivo* NPSCs express high levels of laminin-specific integrin receptors and relatively low levels of collagen receptors ^30,33^. While ECM type is recognized as a determinant of NPSC function, many NPSC EV studies do not specify the type of ECM used. We show here for the first time that certain ECM types, notably collagen IV, may compromise NPSC EV bioactivity, whereas others may enhance bioactivity (i.e., laminin and fibronectin). Thus, we propose ECM type as an important consideration for future NPSC EV studies, with laminin or fibronectin being preferred.

ECM composition is increasingly recognized as a regulator of *in vivo* NPSC function and neurogenesis ^35^, and changes in ECM composition are implicated in neurodegenerative diseases. Neurogenesis in the adult brain occurs in two regions rich in NPSCs: i) the subgranular zone of the dentate gyrus in the hippocampus and ii) the subventricular zone on the lateral ventricles. Here, NPSCs may be quiescent (type B cells) or give rise to type C transit-amplifying cells directly involved in neurogenesis via generation of neuroblasts or OPCs ^36^. A dense and intricate ECM (termed fractones) makes direct contact with NPSCs to promote neurogenesis via integrin signaling and sequestration of neurogenic growth factors ^35,37^. Within this ECM, laminin signaling plays a key role in potentiation of NPSC neurogenic functions and is essential for CNS and PNS axon regeneration ^38-41^. Laminin is a class of heterotrimeric glycoproteins assembled from five alpha, three beta, and three gamma subunits with different isoforms performing distinct neurogenic functions ^42^. Laminin-111, which was used in our study, was the first identified laminin and is frequently utilized for NPSC culture due to its promotion of NPSC proliferation and survival ^32^. However, whether this is the ideal laminin isoform for culturing NPSC for neuroregenerative applications is unclear. Recent studies have shown laminin-211 can enhance oligodendrocyte maturation, and the α5 chain-containing laminin-511 or -521 can enhance NPSC neurogenic functions and may be critical for hippocampal neurogenesis ^43-46^. Another recent approach towards recreating a neurogenic-permissive microenvironment for NPSCs involves the use of laminin-embedded hydrogels to more accurately recapitulate in vivo structural and mechanical cues ^47^. Thus, future studies of optimal laminin subtype and biomaterial matrices hold promise for further enhancing NPSC EV neuroregenerative bioactivity.

We also sought to investigate the role of growth factors in culture on NPSC EV neurite outgrowth bioactivity using a PC-12 cell model. EGF/bFGF are common mitogens included in NPSC culture, whereas NGF is not typically included ^48^. We found that the presence of either EGF/bFGF or NGF is essential for NPSC EV bioactivity, with NGF alone producing more consistently enhanced PC-12 neurite outgrowth compared to EGF/bFGF alone. Recently, engineering NPSCs to overexpress NGF was shown to enhance SCI recovery in mice following transplantation of these cells into the lesion area ^49^. A well-recognized advantage of EVs is their ability to transport and deliver bioactive cargo. Thus, NGF could be leveraged to enhance NPSC EV bioactivity either as a culture media additive as shown here, or by delivering recombinant NGF with EVs either via cargo internalization or surface display techniques.

Neuroinflammation is a pathological hallmark and therapeutic target in neurodegenerative disease and CNS injury. Neuroinflammation is mediated primarily by microglia within the CNS, which are functionally similar to macrophages in the periphery. Here, we show that NPSC EVs also possess anti-inflammatory effects in an LPS-stimulated mouse macrophage assay. However, ECM and growth factors did not have a major impact on NPSC EV anti-inflammatory bioactivity.

The ability of NPSC EVs to increase neurite outgrowth and suppress inflammation suggests promise for treating *in vivo* models of nerve injury. Indeed, we observed NPSC EVs cultured on fibronectin with NGF significantly enhanced axon count and axon diameter and reduced muscle atrophy in a peripheral nerve crush injury rat model. These results suggest that NPSC EVs represent an emerging therapeutic option for enhanced nerve regeneration and muscle recovery following nerve injury.

## 5. Conclusion

We screened commonly-used ECMs for NPSC culture and evaluated NPSC EV bioactivity related to PC-12 neuron proliferation and neurite outgrowth, as well as anti-inflammatory effects. While the conditions selected are only a subset of the overall possible combinations, the data generally support the concept that optimizing the ECM components and growth factors in NPSC culture will be critical for enabling rational design of optimal microenvironments for neurotherapeutic EV production. With further studies in this area, it may eventually be possible to produce EVs with bioactivity specific to what is desired for different applications (e.g., neurogenesis, remyelination, adaptive neural plasticity, reducing neuroinflammation, promoting angiogenesis, etc.)

## 6. Acknowledgements

This research was supported in part by the Maryland Stem Cell Research Fund (2020-MSCRFD-5384 to X.J.; 2020-MSCRFF-5368 to D.D.); the National Institutes of Health (HL141611 and NS110637 to S.M.J.; NS117102 to X.J.) and a University of Maryland MPower Graduate Fellowship (to N.H.P.).

## 8. Data Availability Statement

The data that support the findings of this study are available from the corresponding author upon reasonable request.

## Author Contributions

D.D., N.H.P., and D. L. performed in vitro experiments; X.X. performed in vivo experiments; N.H.P., D.D., D.L., J.T., N.S. X.X. and Z.W. analyzed the data; N.H.P., D.D., X.X. and Z.W. wrote the initial draft of the manuscript; and X.J. and S.M.J. conceived the original idea, designed the experiments, and finalized the manuscript. The authors read and approved the final manuscript.

## Institutional Review Board Statement

The animal study was approved by the Institute Animal Care and Use Committee at the University of Maryland, Baltimore.

---

**Supplemental Table 1.**
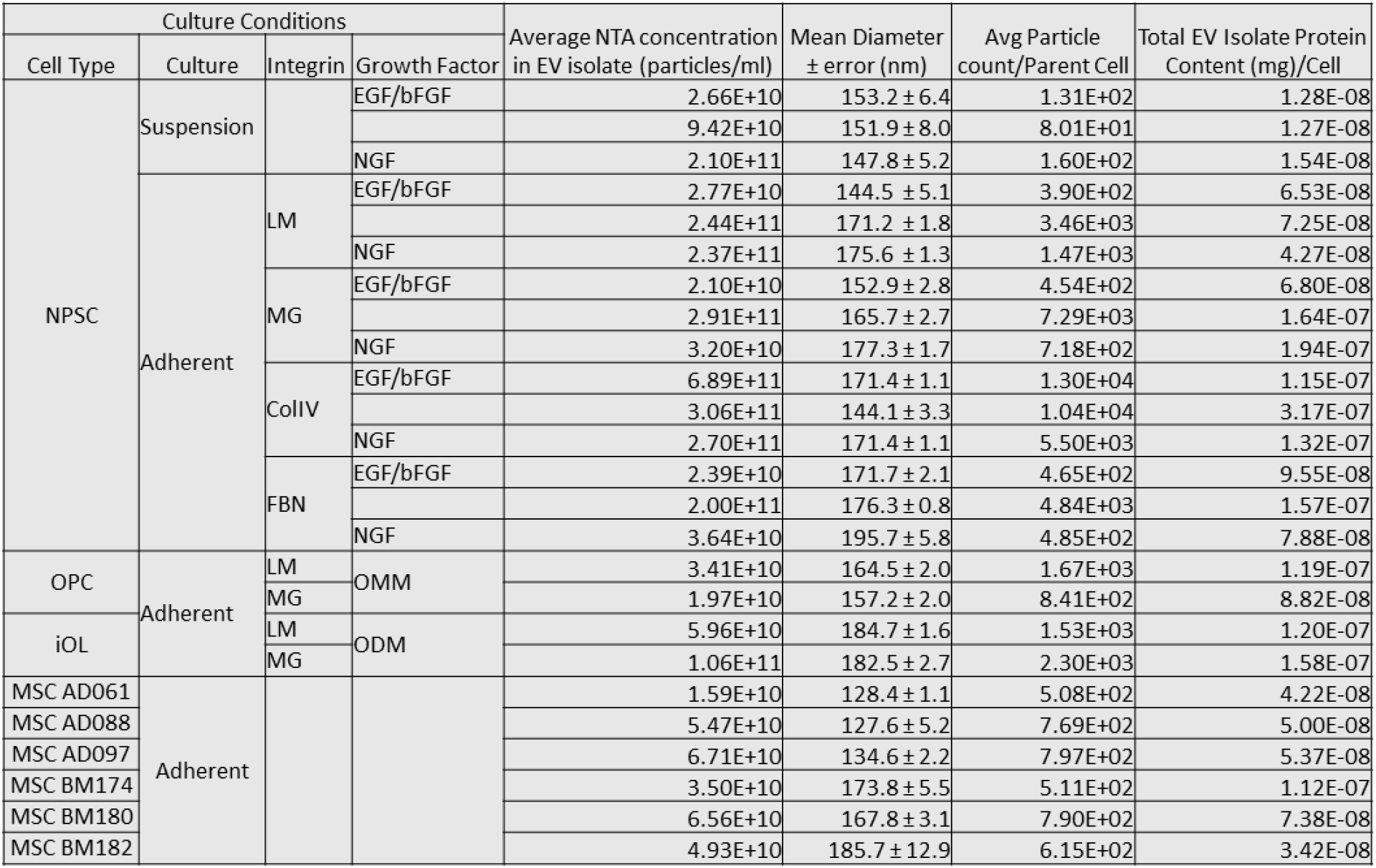
EV characterization and yields. Nanoparticle tracking analysis and bicinchoninic acid assay were used to determine per cell EV yields (particle count and protein content) and EV sample concentrations and mean size across all culture conditions tested.

**Supplemental Figure 1.**
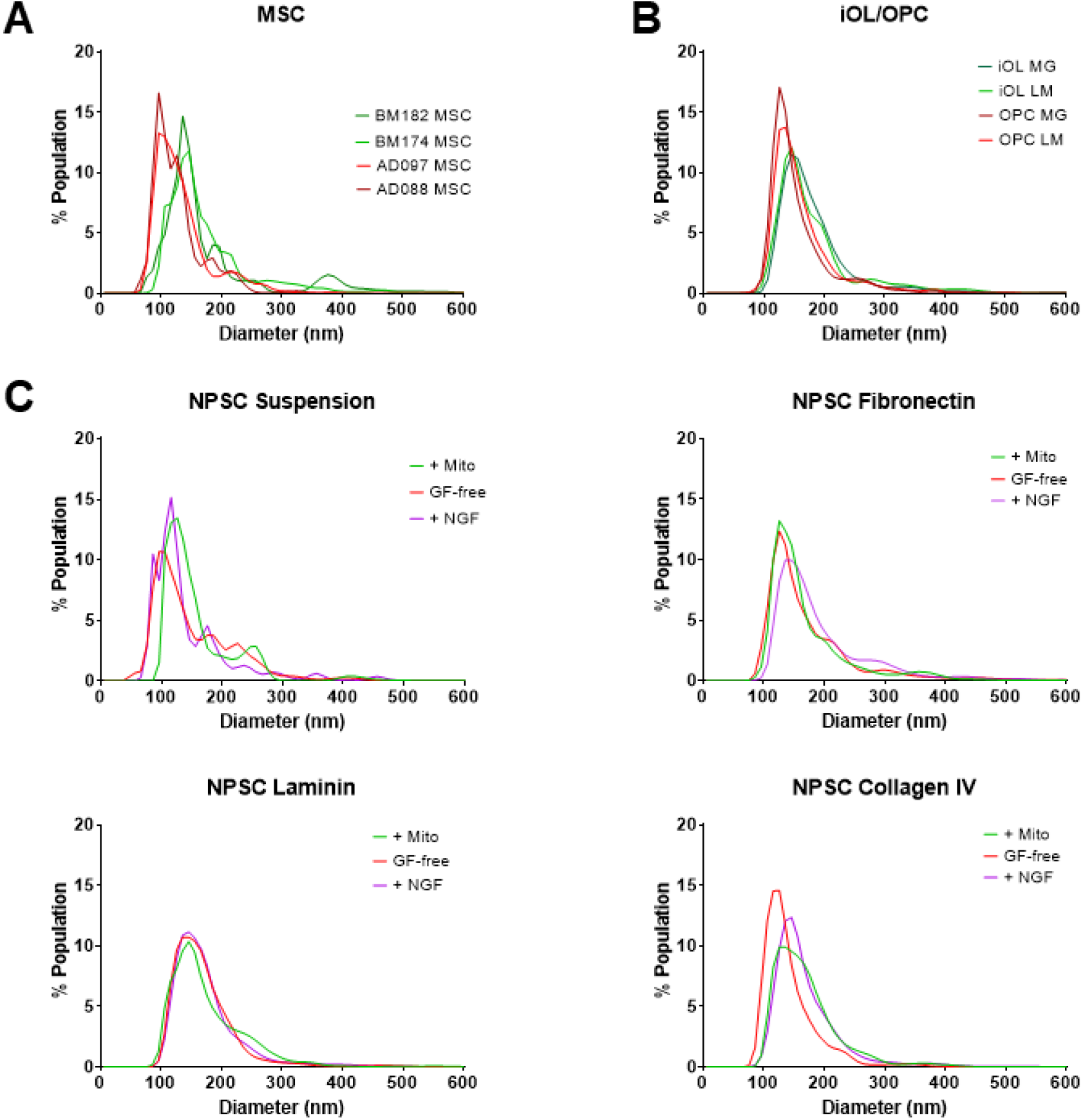
EV characterization. Nanoparticle tracking analysis of EVs from **A)** MSCs from different patient donors and tissue sources (bone marrow or adipose), **B)** iOLs and OPCs grown on Matrigel or laminin, and **C)** NPSCs grown in suspension or on fibronectin, laminin, or collagen IV and in the presence or absence of growth factors (EGF/bFGF) or NGF.

**Supplemental Figure 2.**
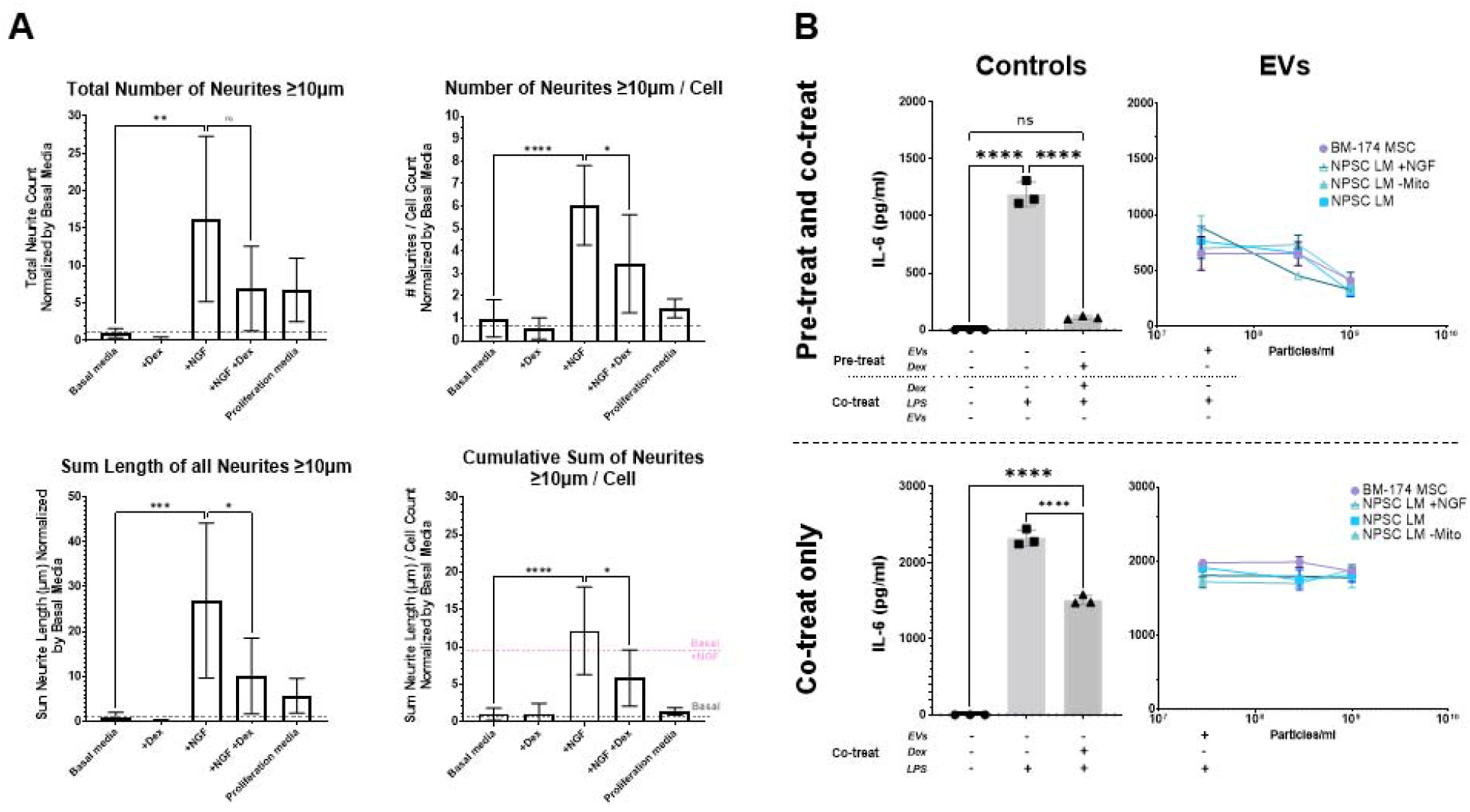
PC-12 neurite outgrowth assay and RAW264.7 immunoassay development. A) Several methods for data analysis were compared and cumulative sum of neurites ≥10μm normalized by cell count was selected for its ability to maximize dynamic range while controlling for cell count. **B)** Pre- and co-treat vs co-treat only regimens were compared and only pre- and co-treat (upper panel) revealed anti-inflammatory effects of EVs.

**Supplemental Figure 3.**
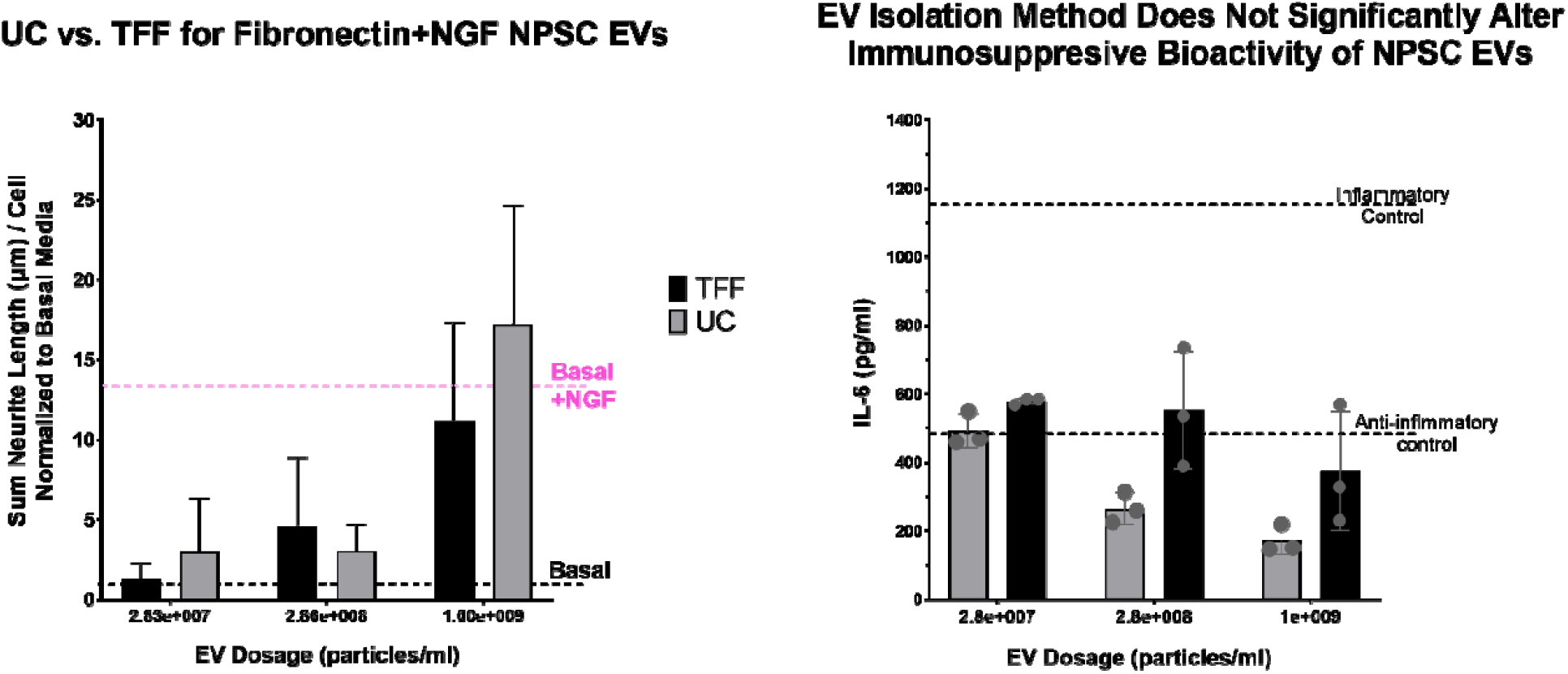
Bioactivity of NPSC EVs is similar between isolation with ultracentrifugation (UC) and tangential flow filtration with 100kDa membrane (TFF). EVs from NPSCs cultured on fibronectin with NGF were evaluated in A) PC-12 neurite outgrowth assay and B) RAW 264.7 mouse macrophage immunoassay.

## References

1 Zhang, B., Yeo, R. W., Tan, K. H. & Lim, S. K. Focus on Extracellular Vesicles: Therapeutic Potential of Stem Cell-Derived Extracellular Vesicles. Int J Mol Sci 17, 174, doi:10.3390/ijms17020174 (2016).

2 Dutta, D., Khan, N., Wu, J. & Jay, S. M. Extracellular Vesicles as an Emerging Frontier in Spinal Cord Injury Pathobiology and Therapy. Trends in Neurosciences, doi:10.1016/j.tins.2021.01.003 (2021).

3 Ramotowski, C., Qu, X. & Villa-Diaz, L. G. Progress in the Use of Induced Pluripotent Stem Cell-Derived Neural Cells for Traumatic Spinal Cord Injuries in Animal Populations: Meta-Analysis and Review. Stem Cells Transl Med 8, 681–693, doi:10.1002/sctm.18-0225 (2019).

4 Webb, R. L. et al. Human Neural Stem Cell Extracellular Vesicles Improve Recovery in a Porcine Model of Ischemic Stroke. Stroke 49, 1248–1256, doi:10.1161/strokeaha.117.020353 (2018).

5 Webb, R. L. et al. Human Neural Stem Cell Extracellular Vesicles Improve Tissue and Functional Recovery in the Murine Thromboembolic Stroke Model. Translational Stroke Research 9, 530–539, doi:10.1007/s12975-017-0599-2 (2018).

6 Patel, D. B., Santoro, M., Born, L. J., Fisher, J. P. & Jay, S. M. Towards rationally designed biomanufacturing of therapeutic extracellular vesicles: impact of the bioproduction microenvironment. Biotechnology Advances 36, 2051–2059, doi:10.1016/j.biotechadv.2018.09.001 (2018).

7 Théry, C. et al. Minimal information for studies of extracellular vesicles 2018 (MISEV2018): a position statement of the International Society for Extracellular Vesicles and update of the MISEV2014 guidelines. Journal of Extracellular Vesicles 7, 1535750, doi:10.1080/20013078.2018.1535750 (2018).

8 Kim, J. B. et al. Oct4-induced oligodendrocyte progenitor cells enhance functional recovery in spinal cord injury model. Embo j 34, 2971–2983, doi:10.15252/embj.201592652 (2015).

9 Sharp, J., Frame, J., Siegenthaler, M., Nistor, G. & Keirstead, H. S. Human embryonic stem cell-derived oligodendrocyte progenitor cell transplants improve recovery after cervical spinal cord injury. Stem Cells 28, 152–163, doi:10.1002/stem.245 (2010).

10 Keirstead, H. S. et al. Human embryonic stem cell-derived oligodendrocyte progenitor cell transplants remyelinate and restore locomotion after spinal cord injury. J Neurosci 25, 4694–4705, doi:10.1523/jneurosci.0311-05.2005 (2005).

11 Barakat, D. J. et al. Survival, integration, and axon growth support of glia transplanted into the chronically contused spinal cord. Cell Transplant 14, 225–240, doi:10.3727/000000005783983106 (2005).

12 Rodriguez, J. P. et al. Abrogation of β-catenin signaling in oligodendrocyte precursor cells reduces glial scarring and promotes axon regeneration after CNS injury. J Neurosci 34, 10285–10297, doi:10.1523/jneurosci.4915-13.2014 (2014).

13 Chen, L. X. et al. Neuroprotective effects of oligodendrocyte progenitor cell transplantation in premature rat brain following hypoxic-ischemic injury. PLoS One 10, e0115997, doi:10.1371/journal.pone.0115997 (2015).

14 Filous, A. R. et al. Entrapment via synaptic-like connections between NG2 proteoglycan+ cells and dystrophic axons in the lesion plays a role in regeneration failure after spinal cord injury. J Neurosci 34, 16369–16384, doi:10.1523/jneurosci.1309-14.2014 (2014).

15 Busch, S. A. et al. Adult NG2+ cells are permissive to neurite outgrowth and stabilize sensory axons during macrophage-induced axonal dieback after spinal cord injury. J Neurosci 30, 255–265, doi:10.1523/jneurosci.3705-09.2010 (2010).

16 Gibson, E. M. et al. Neuronal activity promotes oligodendrogenesis and adaptive myelination in the mammalian brain. Science 344, 1252304, doi:10.1126/science.1252304 (2014).

17 Hamanaka, G., Ohtomo, R., Takase, H., Lok, J. & Arai, K. White-matter repair: Interaction between oligodendrocytes and the neurovascular unit. Brain Circ 4, 118–123, doi:10.4103/bc.bc_15_18 (2018).

18 Lu, P., Jones, L. L., Snyder, E. Y. & Tuszynski, M. H. Neural stem cells constitutively secrete neurotrophic factors and promote extensive host axonal growth after spinal cord injury. Exp Neurol 181, 115–129, doi:10.1016/s0014-4886(03)00037-2 (2003).

19 Barnabé-Heider, F. et al. Origin of new glial cells in intact and injured adult spinal cord. Cell Stem Cell 7, 470–482, doi:10.1016/j.stem.2010.07.014 (2010).

20 Dugas, J. C., Tai, Y. C., Speed, T. P., Ngai, J. & Barres, B. A. Functional genomic analysis of oligodendrocyte differentiation. J Neurosci 26, 10967–10983, doi:10.1523/jneurosci.2572-06.2006 (2006).

21 Fumagalli, M. et al. Phenotypic changes, signaling pathway, and functional correlates of GPR17-expressing neural precursor cells during oligodendrocyte differentiation. J Biol Chem 286, 10593–10604, doi:10.1074/jbc.M110.162867 (2011).

22 Dugas, J. C., Ibrahim, A. & Barres, B. A. The T3-induced gene KLF9 regulates oligodendrocyte differentiation and myelin regeneration. Mol Cell Neurosci 50, 45–57, doi:10.1016/j.mcn.2012.03.007 (2012).

23 Théry, C., Amigorena, S., Raposo, G. & Clayton, A. Isolation and characterization of exosomes from cell culture supernatants and biological fluids. Curr Protoc Cell Biol Chapter 3, Unit 3.22, doi:10.1002/0471143030.cb0322s30 (2006).

24 Schindelin, J. et al. Fiji: an open-source platform for biological-image analysis. Nat Methods 9, 676–682, doi:10.1038/nmeth.2019 (2012).

25 Huang, H., He, W., Tang, T. & Qiu, M. Immunological Markers for Central Nervous System Glia. Neurosci Bull, doi:10.1007/s12264-022-00938-2 (2022).

26 Du, J. et al. Optimal electrical stimulation boosts stem cell therapy in nerve regeneration. Biomaterials 181, 347–359, doi:10.1016/j.biomaterials.2018.07.015 (2018).

27 Xie, Y. et al. Adipose Mesenchymal Stem Cell-Derived Exosomes Enhance PC12 Cell Function through the Activation of the PI3K/AKT Pathway. Stem Cells Int 2021, 2229477, doi:10.1155/2021/2229477 (2021).

28 Kang, H. & Lichtman, J. W. Motor axon regeneration and muscle reinnervation in young adult and aged animals. J Neurosci 33, 19480–19491, doi:10.1523/jneurosci.4067-13.2013 (2013).

29 Li, Y. C., Tsai, L. K., Wang, J. H. & Young, T. H. A neural stem/precursor cell monolayer for neural tissue engineering. Biomaterials 35, 1192–1204, doi:10.1016/j.biomaterials.2013.10.066 (2014).

30 Flanagan, L. A., Rebaza, L. M., Derzic, S., Schwartz, P. H. & Monuki, E. S. Regulation of human neural precursor cells by laminin and integrins. J Neurosci Res 83, 845–856, doi:10.1002/jnr.20778 (2006).

31 Hiraoka, M., Kato, K., Nakaji-Hirabayashi, T. & Iwata, H. Enhanced survival of neural cells embedded in hydrogels composed of collagen and laminin-derived cell adhesive peptide. Bioconjug Chem 20, 976–983, doi:10.1021/bc9000068 (2009).

32 Komura, T., Kato, K., Konagaya, S., Nakaji-Hirabayashi, T. & Iwata, H. Optimization of surface-immobilized extracellular matrices for the proliferation of neural progenitor cells derived from induced pluripotent stem cells. Biotechnology and Bioengineering 112, 2388–2396, doi:https://doi.org/10.1002/bit.25636 (2015).

33 Prowse, A. B., Chong, F., Gray, P. P. & Munro, T. P. Stem cell integrins: implications for ex-vivo culture and cellular therapies. Stem Cell Res 6, 1–12, doi:10.1016/j.scr.2010.09.005 (2011).

34 Frühbeis, C. et al. Oligodendrocytes support axonal transport and maintenance via exosome secretion. PLoS Biol 18, e3000621, doi:10.1371/journal.pbio.3000621 (2020).

35 Kerever, A. et al. Novel extracellular matrix structures in the neural stem cell niche capture the neurogenic factor fibroblast growth factor 2 from the extracellular milieu. Stem Cells 25, 2146–2157, doi:10.1634/stemcells.2007-0082 (2007).

36 Gonzalez-Perez, O. Neural stem cells in the adult human brain. Biol Biomed Rep 2, 59–69 (2012).

37 Mercier, F., Kitasako, J. T. & Hatton, G. I. Anatomy of the brain neurogenic zones revisited: fractones and the fibroblast/macrophage network. J Comp Neurol 451, 170–188, doi:10.1002/cne.10342 (2002).

38 Guldager Kring Rasmussen, D. & Karsdal, M. A. in Biochemistry of Collagens, Laminins and Elastin (ed Morten A. Karsdal) 163–196 (Academic Press, 2016).

39 Aumailley, M. et al. A simplified laminin nomenclature. Matrix Biol 24, 326–332, doi:10.1016/j.matbio.2005.05.006 (2005).

40 Grimpe, B. et al. The critical role of basement membrane-independent laminin gamma 1 chain during axon regeneration in the CNS. J Neurosci 22, 3144–3160, doi:10.1523/jneurosci.22-08-03144.2002 (2002).

41 Kazanis, I. et al. Quiescence and activation of stem and precursor cell populations in the subependymal zone of the mammalian brain are associated with distinct cellular and extracellular matrix signals. J Neurosci 30, 9771–9781, doi:10.1523/jneurosci.0700-10.2010 (2010).

42 Kazanis, I. & ffrench-Constant, C. Extracellular matrix and the neural stem cell niche. Dev Neurobiol 71, 1006–1017, doi:10.1002/dneu.20970 (2011).

43 Chun, S. J., Rasband, M. N., Sidman, R. L., Habib, A. A. & Vartanian, T. Integrin-linked kinase is required for laminin-2-induced oligodendrocyte cell spreading and CNS myelination. J Cell Biol 163, 397–408, doi:10.1083/jcb.200304154 (2003).

44 Dunville, K. et al. Laminin 511 and WNT signalling sustain prolonged expansion of hiPSC-derived hippocampal progenitors. Development 149, doi:10.1242/dev.200353 (2022).

45 Indyk, J. A., Chen, Z. L., Tsirka, S. E. & Strickland, S. Laminin chain expression suggests that laminin-10 is a major isoform in the mouse hippocampus and is degraded by the tissue plasminogen activator/plasmin protease cascade during excitotoxic injury. Neuroscience 116, 359–371, doi:10.1016/s0306-4522(02)00704-2 (2003).

46 Hyysalo, A. et al. Laminin α5 substrates promote survival, network formation and functional development of human pluripotent stem cell-derived neurons in vitro. Stem Cell Res 24, 118–127, doi:10.1016/j.scr.2017.09.002 (2017).

47 Barros, D., Amaral, I. F. & Pêgo, A. P. Laminin-Inspired Cell-Instructive Microenvironments for Neural Stem Cells. Biomacromolecules 21, 276–293, doi:10.1021/acs.biomac.9b01319 (2020).

48 Ahmed, A. K., Isaksen, T. J. & Yamashita, T. Protocol for mouse adult neural stem cell isolation and culture. STAR protocols 2, 100522 (2021).

49 Wang, L. et al. Neural Stem Cells Overexpressing Nerve Growth Factor Improve Functional Recovery in Rats Following Spinal Cord Injury via Modulating Microenvironment and Enhancing Endogenous Neurogenesis. Frontiers in Cellular Neuroscience 15, doi:10.3389/fncel.2021.773375 (2021).

